# Cold tolerance and diapause within and across trophic levels: endoparasitic wasps and their fly host have similar phenotypes

**DOI:** 10.1101/2023.01.04.522725

**Authors:** Trinity McIntyre, Lalitya Andaloori, Glen Ray Hood, Jeffrey L. Feder, Daniel A. Hahn, Gregory J. Ragland, Jantina Toxopeus

## Abstract

Low temperatures associated with winter can limit the survival of organisms, especially ectotherms whose body temperature is similar to their environment. Important adaptations for overwintering such as cold hardiness and diapause have been well-explored in many insect taxa. However, there is a gap in understanding how overwintering may vary among groups of species that interact closely, such as multiple parasitoid species that attack the same host insect. Our study investigated cold tolerance and diapause phenotypes in three endoparasitoid wasps of the apple maggot fly *Rhagoletis pomonella* (Diptera: Tephritidae): *Utetes canaliculatus, Diachasma alloeum*, and *Diachasmimorpha mellea* (Hymenoptera: Braconidae). Using a combination of respirometry and eclosion tracking, we detected diapause phenotypes in all three wasp species, remarkably similar to the fly host. Weak diapause was rare (< 5%) in all three wasp species, and while most *D. mellea* (93%) entered prolonged diapause under warm conditions, the majority of *U. canaliculatus* (92%) and *D. alloeum* (72%) averted diapause (non-diapause). There was limited interspecific variation in acute cold tolerance among the three wasp species: wasps and flies had similarly high survival (>87%) following exposure to extreme low temperatures (- 20°C) as long as their body fluids did not freeze. The wasp species showed little interspecific variation in survival following prolonged exposure to mild chilling of 8 or more weeks at 4°C. This study shows remarkable conservation of cold tolerance and diapause phenotypes within and across trophic levels. The interaction between diapause phenotype and cold hardiness in these parasitoids is an interesting direction for future research.

**Graphical Abstract and Highlights:** 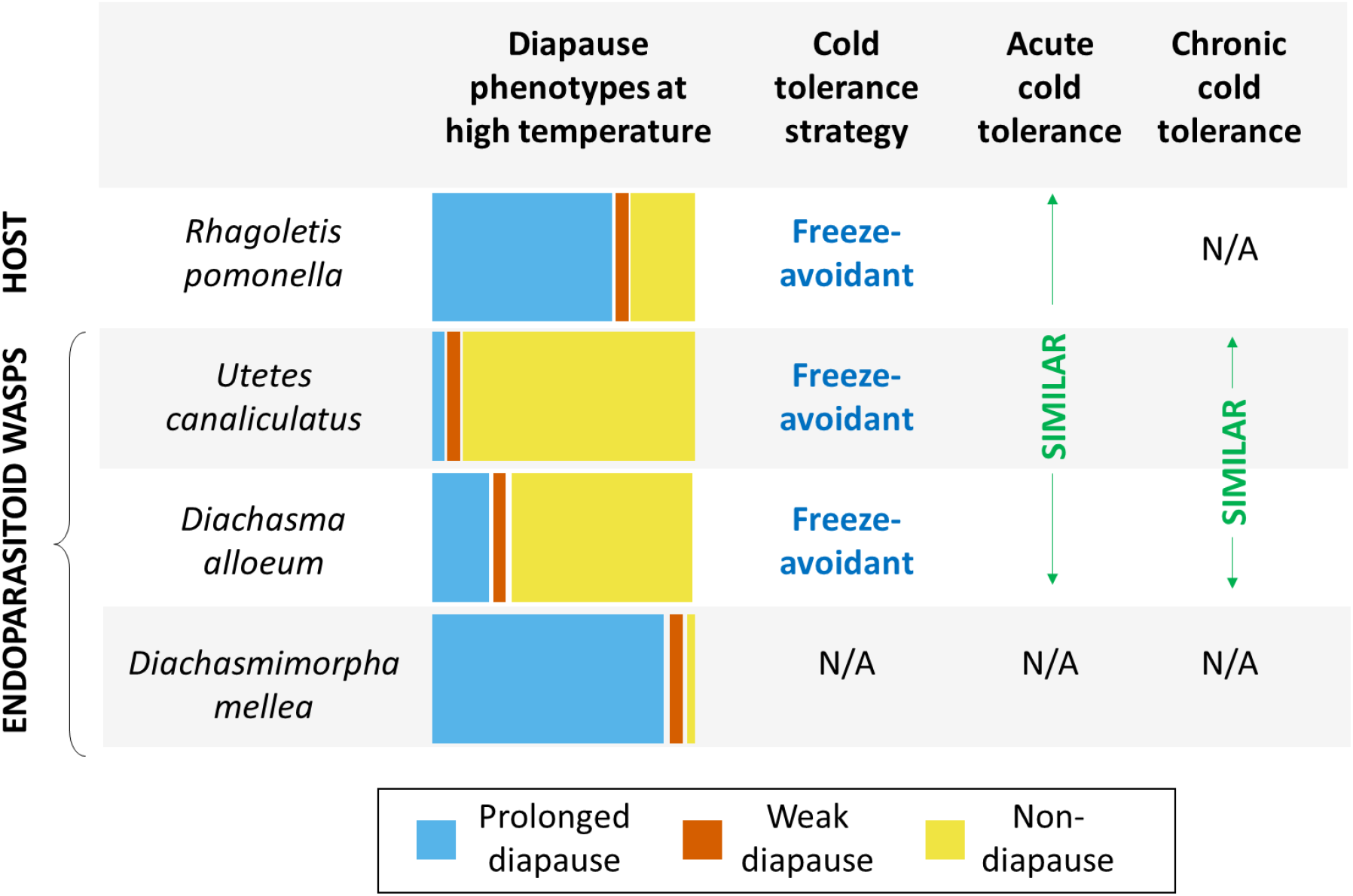

- The apple maggot fly and its parasitoids exhibit the same three diapause phenotypes
- Each parasitoid wasp species exhibits different proportions of these phenotypes
- *Utetes canaliculatus* and *Diachasma alloeum* are freeze-avoidant, like their host fly
- These wasps and flies survive to similarly extreme low temperatures (c. -20°C)
- Each wasp species survives prolonged exposure to mild chilling (4°C) similarly well

## 1. Introduction

Low temperatures associated with winter can impose strong selective pressure on organisms (Marshall et al., 2020; Sinclair et al., 2003; Williams et al., 2015). Ectothermic animals are particularly susceptible to the stressors associated with winter because their internal body temperature generally reflects that of their environment. Temperatures below 0°C can therefore cause ectothermic animal body fluids to freeze, resulting in damage that is often lethal (Lee, 2010; Toxopeus and Sinclair, 2018). Many studies have documented overwintering biology of various insect species, including adaptations such as cold tolerance strategies and diapause (Hand et al., 2016; Lee, 2010; Sinclair et al., 2015; Toxopeus and Sinclair, 2018; Wilsterman et al., 2021). In addition, there is increasing interest in exploring overwintering biology of closely interacting species. For example, the relationship between the cold tolerance and diapause phenotypes of parasitoids and their host species may alter interspecific dynamics as climates change or geographic ranges of populations shift (Colinet and Boivin, 2011; Le Lann et al., 2021). It is in this vein that we investigate cold tolerance and diapause in a guild of interacting endoparasitoid wasps and their host, the apple maggot fly, *Rhagoletis pomonella* Walsh (Diptera: Tephritidae).

Insects that overwinter in temperate climates have evolved physiological cold tolerance strategies to survive the challenges associated with low temperatures. Freeze-tolerant insects survive freezing of their body fluids, while freeze-avoidant insects depress the temperature at which ice formation begins (supercooling point; SCP) to survive sub-zero temperatures in an unfrozen state (Lee, 2010; Toxopeus and Sinclair, 2018). The ability to survive short exposures to extreme temperatures is important for defining an insect’s lower lethal temperature (Sinclair et al., 2015; Toxopeus et al., 2019, 2016). However, many temperate insects overwinter in relatively mild environments (e.g., temperatures close to 0°C in the soil under a layer of insulative snow cover). Therefore, characterizing survival following both short exposure to extreme conditions (acute cold tolerance) and milder prolonged chilling (chronic cold tolerance) is important for predicting overwintering survival (Roberts et al., 2021; Sinclair et al., 2015).

Diapause is another important adaptation for insects overwintering in temperate climates. Prior to the onset of winter, many temperate insects enter diapause, a state of developmentally-programmed dormancy associated with metabolic rate suppression and enhanced stress tolerance (Hand et al., 2016; Koštál, 2006; Wilsterman et al., 2021). While diapause is obligate (necessary for completion of the life cycle) in some insect species, many species exhibit interindividual variation in diapause phenotypes based on environmental conditions (Hodek and Hodková, 1988; Koštál, 2006). For example, relatively warm fall temperatures may cause some individuals of a diapause-capable species to completely forgo (avert) diapause prior to winter (Hodek and Hodková, 1988; Koštál, 2006). In this study, we refer to these as ‘non-diapause’ phenotypes (Calvert et al., 2022; Dambroski and Feder, 2007; Toxopeus et al., 2021). In mild winter climates, these non-diapause insects may be winter active (Alfaro-Tapia et al., 2021; Tougeron et al., 2017). Exposure to warm conditions prior to winter can also cause insects to remain in diapause for only a short duration, a phenotype which is commonly referred to as ‘weak diapause’ (Toxopeus et al., 2021) or ‘shallow diapause’ (Calvert et al., 2022; Dambroski and Feder, 2007). This weak or shallow diapause phenotype has not been documented in many insects (Masaki, 2002; Wilsterman et al., 2021), and can be contrasted with animals in ‘prolonged diapause’. In both weak and prolonged diapause phenotypes, insects enter into a programmed dormancy (Wilsterman et al., 2021). However, individuals in prolonged diapause are more recalcitrant in responding to the cues that drive diapause termination (e.g., warm temperatures) than individuals in weak diapause (Wilsterman et al., 2021). Populations containing a mix of weak and prolonged diapause individuals can exhibit bi- or multi-modal distributions of diapause termination and resumption of post-diapause development over time (Calvert et al., 2022; Dambroski and Feder, 2007; Masaki, 2002). Diapause is often necessary for overwintering survival and for the proper synchronization of growth and reproduction with favorable environments (Hand et al., 2016; Koštál, 2006; Tauber and Tauber, 1976). Thus, averting or prematurely terminating diapause can cause decreased cold tolerance and overwintering survival (Boiteau and Coleman, 1996; Ciancio et al., 2021; Lee and Denlinger, 1985; Lehmann et al., 2018; Toxopeus et al., 2021).

Several studies have examined the cold tolerance and diapause of parasitoids, often with a focus on their potential as biocontrol agents (Colinet and Boivin, 2011; Le Lann et al., 2021). Endoparasitoids feed and develop to adulthood within a single host organism, usually killing the host in the process (Godfray, 1994; Hood et al., 2021). The parasitoid cold tolerance literature is dominated by studies of wasps (Hymenoptera) that parasitize insect pests from many different insect orders, including Diptera (e.g., drosophilids; Amiresmaeili et al., 2020; Li et al., 2015; Murata et al., 2013), Lepidoptera (e.g., pyralid moths; Carrillo et al., 2005; Foray et al., 2013), Coleoptera (e.g., emerald ash borer; Hanson et al., 2013), and Hemiptera (e.g., aphids; Alford et al., 2017; Colinet and Hance, 2010; Tougeron et al., 2018). However, studies that examine multiple parasitoids that attack a single host are rare (e.g., Hanson et al., 2013), and in these cases the immature life stages of multiple parasitoid species may never interact due to distinct geographic distributions (e.g., Murata et al., 2013) or strong differences in life history timing (e.g., Le Lann et al., 2011). *Rhagoletis pomonella* is a pest of commercial apples that is parasitized by multiple wasp species with overlapping geographic distributions and life history timing (Feder, 1995; Hood et al., 2012). This system provides an opportunity to study the cold hardiness and diapause of species both within a guild of parasitoids and across trophic levels.

Genetic, evolutionary, and physiological studies in *R. pomonella* have established that overwintering, diapausing pupae have relatively low and invariant SCPs, but their diapause development can be disrupted by high temperatures (Calvert et al., 2022; Dambroski and Feder, 2007; Toxopeus et al., 2021). Adult *R. pomonella* lay their eggs in ripe fruit of hawthorns (*Crataegus* spp.) and introduced, domesticated apples (*Malus domestica*) in many regions of North America during the late summer or fall (Dean and Chapman, 1973; Feder et al., 1993; Hood et al., 2013). Larvae develop within those fruit, burrow out of fruit that have dropped to the ground, and then pupate in the soil, where they remain for 10 to 11 months of the year (Boller and Prokopy, 1976; Dean and Chapman, 1973). Most of these pupae in the northern part of the flies’ range enter a prolonged diapause and remain recalcitrant to diapause termination and post-diapause development until late summer of the following year (Calvert et al., 2022; Dambroski and Feder, 2007; Toxopeus et al., 2021). Overwintering *R. pomonella* pupae from apple fruits are freeze-avoidant down to c. -20°C and have similar acute cold tolerance regardless of diapause status, but only diapausing pupae survive prolonged chilling well (> 50% survival following 24 weeks at 4°C; Toxopeus et al., 2021). Although prolonged diapause is likely important for overwintering survival in natural environments, both non-diapause and weak diapause pupae have been observed when *R. pomonella* are held in warm (>20°C) conditions for several weeks in the laboratory (Calvert et al., 2022; Dambroski and Feder, 2007; Toxopeus et al., 2021). In some populations, more than half of the individuals may avert or prematurely terminate diapause under these conditions, and the proportion that do so can vary with latitude and host fruit (Calvert et al., 2022; Dambroski and Feder, 2007; Toxopeus et al., 2021). If exposed to simulated winter conditions, non-diapause and weak diapause individuals typically eclose as adults more quickly post-winter than prolonged diapause individuals (Calvert et al., 2022; Dambroski and Feder, 2007; Toxopeus et al., 2021). Combined studies applying artificial selection and population genetic surveys suggest that natural selection is acting on these diapause phenotypes in populations that experience relatively longer and warmer pre-winter conditions (Doellman et al., 2019; Dowle et al., 2020; Egan et al., 2015; Hood et al., 2020; Powell et al., 2020; Ragland et al., 2017), with potential implications for their overwintering survival.

Three endoparasitoid wasps (Hymenoptera: Braconidae: Opiinae) attack *R. pomonella* in the Northeastern and midwestern United States: *Utetes canaliculatus* Gahan, *Diachasma alloeum* (Muesebeck), and *Diachasmimorpha mellea* Gahan (Forbes et al., 2010; Hood et al., 2015, 2012; Wharton and Marsh, 1978). In these regions, *D. alloeum* is usually the most prevalent parasitoid of *R. pomonella* that infest apples and hawthorn fruits, followed by *U. canaliculatus* and *D. mellea* (Feder, 1995; Hood et al., 2015, 2012). Each parasitoid’s life history is closely linked to that of its host but varies slightly among species. *Utetes canaliculatus* oviposits into *R. pomonella* eggs, and has an earlier but overlapping eclosion phenology with *D. mellea* and the later emerging *D. alloeum*, which each oviposit into second and third instar larvae of *R. pomonella* (Forbes et al., 2010; Hood et al., 2015). Although multiple species may infest the same immature fly (Hood et al., 2012), only one wasp larva will survive to completely consume the host within its puparium (tanned outer case) within 10 to 15 d post-pupariation (Forbes et al., 2010; Hood et al., 2015; Lathrop and Newton, 1933). While most of these wasp larvae are assumed to overwinter in diapause (Forbes et al., 2009; Hood et al., 2015), the diapause phenotypes and ability to survive extreme low temperature of each parasitoid species have not been systematically characterized in the same detail as their fly host (cf. Calvert et al., 2022; Dambroski and Feder, 2007; Toxopeus et al., 2021).

Our goal was to determine whether the guild of endoparasitoid wasp species that attack *R. pomonella* fruit flies exhibit similar or different overwintering cold tolerance compared to their host, and whether the parasitoids might also be susceptible to diapause disruption under warm pre-winter conditions. Although many *R. pomonella* studies compare populations that infest apple and hawthorn fruits (e.g., Calvert et al., 2022; Doellman et al., 2019; Dowle et al., 2020; Egan et al., 2015; Hood et al., 2020; Powell et al., 2020; Ragland et al., 2017), here we focus on hawthorn-infesting populations only, which tend to have higher rates of parasitism from each of the three wasp species than populations attacking apple-infesting flies (Feder, 1995; Hood, 2016; Hood et al., 2015). We characterize and compare the cold tolerance of *U. canaliculatus, D. alloeum*, and *D. mellea* to each other and to *R. pomonella* collected from hawthorn fruits using several metrics, including cold tolerance strategy, SCP, acute cold tolerance, and chronic cold tolerance. In addition, we use respirometry and laboratory eclosion experiments of the wasps to detect variation in diapause phenotypes for comparison to their fly host. Given increasingly warmer fall and winter temperatures (Le Lann et al., 2021; Marshall et al., 2020), this study lays the ground work for future investigations of the interaction between diapause phenotype and cold tolerance in these parasitoid species.

## 2. Materials and Methods

### 2.1 Insect collection

We collected *R. pomonella* from infested hawthorn fruit (*Crataegus* spp.) at two sites previously documented to have high rates of parasitism. To characterize diapause phenotypes in each wasp species, we collected fruits from the ground beneath infested trees in September 2009 from Grant, MI, USA (43°21’N 85°51’W). Within 48 h of collection, we transported the fruits at ambient temperature to the University of Notre Dame and University of Florida. As described by Toxopeus et al. (2021), fruits were suspended over plastic trays in mesh wire baskets and held in an environmentally-controlled room at 21 – 22°C. Over a 2-week period, newly formed puparia were collected once daily from the plastic trays and transferred into Petri dishes that were then placed in a Percival DR-36VL incubator (Percival Scientific, Perry, IA) set to 21°C at 14 L:10 D, and 80% relative humidity (R.H.). For the diapause phenotyping described below (section 2.2), pupae were kept at these warm conditions (21°C) throughout the duration of the experiments.

To characterize cold tolerance and post-winter development, we collected fruits in September 2019 in a similar manner as above, but our collection site was at a nearby location in East Lansing, MI, USA (42°42’N 84°29’W) with similarly high parasitism rates. Fruits were transported to and processed at the University of Colorado, Denver, to collect pupae as described above, except that newly formed puparia were transferred to an incubator at a higher temperature (25°C). Pupae were kept in these warm ‘pre-winter’ conditions (25°C) for 10 d post-pupariation, and were then transferred to 4°C in total darkness with c. 85% R.H. to simulate overwintering for 8, 16, and 24 weeks prior to measurement of the cold tolerance metrics described below (sections 2.3 – 2.5).

Although fly pupae and wasp larvae are morphologically distinct, the puparia of *R. pomonella* are opaque, which prevented us from determining whether any given fly was parasitized without removing part or all of the puparium. In addition, larvae of the three wasp species are morphologically indistinguishable from each other. Therefore, most experiments were conducted on unidentified individuals within the intact fly puparia and were identified to the species level after experiments were performed (section 2.6). As a result, sample sizes associated with each of the three parasitoid species were largely determined by the parasitism rates of each of the three species in the populations from which they were collected.

### 2.2. Diapause phenotypes

To determine the prevalence of each of the three diapause phenotypes (prolonged diapause, weak diapause, non-diapause) in *U. canaliculatus, D. alloeum*, and *D. mellea*, we pooled wasp samples from two parallel experiments conducted in 2009. We used 129 wasp-infested fly pupae at the University of Florida to categorize diapause phenotypes via respirometry (details below). We also tracked eclosion of 247 wasps at the University of Notre Dame to categorize diapause phenotypes (details below). While we focus on the diapause phenotypes of wasp species in this study, diapause phenotype incidence in *R. pomonella* from these same populations was also recorded and is reported by Calvert et al. (2022).

The three diapause phenotypes can be distinguished in non-overwintered *R. pomonella* by different patterns of metabolic rate over time at warm temperatures (Ragland et al. 2009, Toxopeus et al. 2021, Calvert et al. 2022), and we found similar patterns in the wasps that could be used to distinguish diapause phenotypes of each parasitoid species. To distinguish these phenotypes, we used stop-flow respirometry at 21°C to measure the rate of CO_2_ production (*V*CO_2_) as a proxy for metabolic rate as previously described in (Ragland et al., 2009). In brief, we measured metabolic rate 5 d post-pupariation and once weekly thereafter for 10 weeks. For each measurement, we first transferred each individual to an airtight syringe and purged the syringe with CO_2_-free air. We then measured the amount of CO_2_ produced in the syringe after 3 h incubation at 21°C using a LiCor 7000 infrared CO_2_ analyzer (Lincoln, NE) interfaced to Sable Systems International Expedata logging software (Las Vegas, NV) at room temperature. Pupae were returned to the 21°C incubator between weekly measurements. We calculated *V*CO_2_ at each time point using manual bolus integration (Lighton, 2018) normalized to control syringes that contained the same CO_2_-free air but no pupae (Ragland et al., 2009).

Similar to their fly host (Calvert et al., 2022), individuals that maintained a high metabolic rate over the first three measurements were classified as non-diapause; these wasps typically eclosed as adults before any additional *V*CO_2_ measurements could be made. In contrast, individuals that decreased their metabolic rate over the first three measurements were classified as either weak diapause or prolonged diapause phenotypes (Calvert et al., 2022). Prolonged diapause individuals maintained a low metabolic rate for the entire 10 weeks of measurement, whereas weak diapause individuals exhibited an increase in metabolic rate (which indicates the end of diapause; Powell et al., 2020; Ragland et al., 2009) between 40 and 75 d post-pupariation and eclosed as adults 2 to 3 weeks after this increase in metabolic rate. Eclosed and uneclosed wasps were stored at -80°C until they could be identified to the species level (section 2.6).

The three diapause phenotypes can also be distinguished in non-overwintered *R. pomonella* by different patterns of eclosion time at warm temperatures (Calvert et al., 2022; Dambroski and Feder, 2007), and we were able to use similar principles to determine wasp diapause phenotype. Individuals were held at 21°C for 100 d in Petri dishes, and eclosion of adults was tracked once daily. Similar to *R. pomonella* (Calvert et al., 2022; Dambroski and Feder, 2007), each wasp species exhibited a multimodal distribution of eclosion phenology post-pupariation, with non-diapause individuals eclosing before weak diapause individuals (Fig. S1). Similar to Dambroski and Feder (2007), we classified individuals as non-diapause if they eclosed as adults in 35 d or less post-pupariation, weak diapause if they eclosed between 36 and 100 d post-pupariation, and prolonged diapause if they failed to eclose prior to 100 d at 21°C (Fig. S1). Eclosed and uneclosed wasps were stored at -80°C until they could be identified to the species level (section 2.6).

The incidence of each diapause phenotype was similar between the respirometry and eclosion time experiments in 2009 (Table S1). We therefore pooled diapause phenotype data from both experiments for statistical analysis and graphical presentation. We compared the proportion of wasps exhibiting each diapause phenotype among species using a G-test (McDonald, 2014). All statistical analyses were conducted in R v4.1.2 (R Core Team, 2022) and the code is available in the supplementary material.

### 2.3 Cold tolerance strategy

To determine cold tolerance strategy, we used the 2019 collection to assess survivorship of frozen and unfrozen (supercooled) wasp larvae and fly pupae exposed to the same low temperature, as described below (Sinclair et al., 2015; Toxopeus et al., 2021). A species was classified as freeze-tolerant if most (>75%) frozen individuals survived, freeze-avoidant if most (>75%) supercooled individuals survived but all frozen individuals died, and chill-susceptible if mortality was high for both supercooled and frozen individuals (Sinclair et al., 2015). The insects were chilled for either 8 or 16 weeks at 4°C prior to these experiments to detect potential differences in cold tolerance over time, e.g. due to acclimation (cf. Toxopeus et al., 2019). Because cold tolerance strategy (a qualitative category) did not vary with chilling duration, we pooled the data from groups of flies and wasps sampled after 8 and 16 weeks of chilling for graphical presentation.

To differentiate between flies and wasps after chilling and prior to the cold tolerance strategy experiments, we uncapped a small anterior portion of the puparium to morphologically identify the insects as either *R. pomonella* or a parasitoid wasp and confirm that individuals were alive prior to this experiment. This procedure does not affect the supercooling point (SCP) or short term (1 week) survival after acute cold exposure for *R. pomonella* (Toxopeus et al., 2021). At this time, any individuals that showed obvious signs of decay or decomposition during uncapping were excluded from the experiments, resulting in sample sizes of 40 wasp larvae and 40 fly pupae, 20 of each following 8 and 16 weeks chilling.

To expose groups of 20 wasps or flies to low temperatures, they were cooled in a custom aluminum block connected to a recirculating Arctic A40 recirculating chiller (ThermoFisher Scientific, Waltham, MA, USA) containing 50% (v/v) propylene glycol (Toxopeus et al., 2021). Each wasp or fly was placed in a separate 1.7 ml microcentrifuge tube. We then cooled each of the groups at -0.25°C min^-1^ from 4°C to a temperature (c. -20°C) at which half of the individuals froze and the other half remained supercooled (Sinclair et al., 2015). To detect freezing (an exothermic process), we measured the temperature of each individual once per second with a type T thermocouple interfaced to PicoLog temperature logging software via a TC-08 interface (Omega Engineering, Norwalk, CT, USA). Following the cooling procedure, individual wasp larvae and fly pupae were immediately transferred into 0.2 ml microcentrifuge tubes and held at 4°C (complete darkness, 85% R.H.) for 1 d, followed by 25°C (14 L:10D, 80% R.H.) for 6 d to allow for recovery (Toxopeus et al., 2021). Survival was assessed after this 1-week recovery period using stop-flow respirometry and visual markers of damage and decay (Ragland et al., 2009; Toxopeus et al., 2021). All wasp larvae were then flash-frozen in liquid nitrogen and stored at -80°C until subsequent genetic analysis to identify wasp species (section 2.6).

### 2.4 Acute cold tolerance

Supercooling point (SCP) is a suitable metric for comparing acute cold tolerance of freeze-avoidant species (Sinclair et al., 2015) because a lower SCP allows an individual to remain unfrozen, and therefore survive, at lower temperatures. Similar to the cold tolerance strategy experiments, we first uncapped pupae to differentiate between flies and wasps and determine that individuals were alive prior to this experiment (no signs of decay or decomposition). We measured SCP after 8 and 16 weeks of chilling in 40 wasp larvae and 40 fly pupae (20 of each per chilling duration). We used similar cooling methods as for the cold tolerance strategy determination described above, except that groups were cooled to a temperature at which all individuals froze (c. -22°C). All frozen wasp larvae were then transferred to -80°C and stored until subsequent genetic analysis to identify wasp species (section 2.6).

Supercooling points were compared among all three wasp species and their fly host. Both mass and mild low temperature acclimation can impact SCP (Sinclair et al., 2015), but we were primarily interested in interspecific differences in SCP. We therefore compared SCP among the four species (*R. pomonella* and each of the three species of endoparasitoids) using an ANCOVA with ‘species’ as a categorical factor and ‘mass’ and ‘chill duration’ (i.e. acclimation time) as fixed effect covariates.

### 2.5 Chronic cold tolerance and post-chill developmental time

To compare tolerance of prolonged, mild chilling among wasp species, we assessed the survival and development of individuals after short (8 weeks), moderate (16 weeks), or prolonged (24 weeks) chilling at 4°C in complete darkness at 85% R.H. In this experiment, chilling began 10 d post-pupariation (see section 2.1), so all individuals experienced a ‘pre-winter’ period of 25°C for 10 d. For each chilling duration, we used 325 individuals, which included an unknown mixture of parasitized and unparasitized fly pupae. Post-chilling, we transferred the Petri dishes containing pupae to 25°C (14 L:10D, 80% R.H.) and checked once weekly for adult eclosion (as a metric of survival) for 100 d. We tracked *R. pomonella* eclosion (see supplementary information), but excluded the flies from our statistical analysis because eclosion is not a complete measure of chronic cold tolerance in this species: after short chill times (e.g., 8 weeks), many flies are still alive but do not eclose as adults because they have not yet completed diapause (cf. Feder et al., 1997; Toxopeus et al., 2021).

To compare chronic cold tolerance among *U. canaliculatus, D. alloeum*, and *D. mellea*, we assessed the relative abundance of each wasp species that eclosed after chill durations of 8, 16, and 24 weeks. We do not have complete survivorship data for each species because many uneclosed (dead) individuals were not identified to the species level; DNA quality degraded substantially post-mortem, preventing genetic identification. Therefore, we analyzed survivorship by focusing only on wasps that eclosed. If all three wasp species had similar chronic cold tolerance, we would expect the species composition (proportion of *D. alloeum, D. mellea*, and *U. canaliculatus*) of eclosing wasps after 16 and 24 weeks chilling to be similar to the species composition after the shortest chilling treatment of 8 weeks. Conversely, if one wasp species was more cold-tolerant than the other species, we would expect the relative abundance of this species in the population of eclosed wasps to increase with prolonged chill times. We used a G-test (McDonald, 2014) to determine whether the species composition of wasps that eclosed after 16 and 24 weeks of chilling (‘observed’ values) was similar to the species composition of each wasps that eclosed after 8 weeks of chilling (‘expected’ values).

### 2.6 Wasp species identification

All eclosed adult wasps were morphologically identified to the species level using diagnostic characteristics provided by the key of Wharton and Yoder (http://paroffit.org). Unclear cases of species identification, including wasps that did not complete development to the adult stage, were genetically identified using a PCR-based assay developed and described by Hood (2016). Briefly, we extracted DNA from each individual using a PureGene DNA extraction kit (Qiagen Inc., Germantown MD, USA). Each individual was homogenized in 100 μl PureGene Cell Lysis solution with a plastic micropestle (Fisher Scientific, Waltham MA, USA), and all other steps followed manufacturer’s instructions, with reagent volumes scaled down to align with the volume of lysis solution. DNA was eluted in 30 μl nuclease-free water. We then used PCR to amplify diagnostic regions of the mitochondrial *cytochrome oxidase subunit I* (*COI*) gene (Forbes et al., 2009; Hood et al., 2015) with species-specific primers that yielded amplicons of different sizes for each species, as described in Hood (2016; Table 1). We performed PCR reactions with DreamTaq (Fisher Scientific, Mississauga, ON, Canada) according to the manufacturer’s instructions in a total volume of 25 μL containing 1 μL of template DNA and used the following thermocycler protocol. After an initial denaturing period of 5 min at 94°C, DNA was amplified for 35 cycles with denaturation at 94°C for 30 s, reannealing for 1 min at the temperature specified in Table 1, and extension at 72°C for 1 min, followed by one final extension for 10 minutes at 72°C. The PCR products from each of the three species-specific primer pairs for a given individual were then pooled and run together in a 1.5% agarose gel for band visualization and sizing compared to a 100 bp ladder; see Fig. S2 for an example gel. Positive controls (DNA from morphologically identified adults) were included in each set of PCR performed, and the sequences of one positive control amplicon per species were confirmed by Sanger Sequencing at The Centre for Applied Genomics (Sick Kids Hospital, Toronto, ON). The taxonomic identity of these sequences was confirmed by using Clustal Omega to align them to known *COI* sequences of the three wasps from NCBI (Figs. S3 – S5).

**Table 1.**
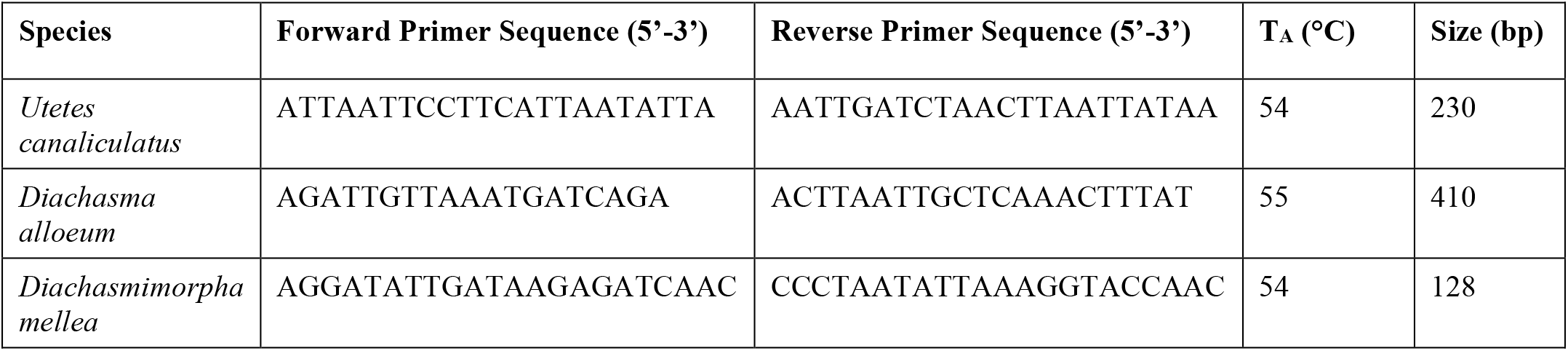
Primers developed by Hood (2016) used to identify three endoparasitoid wasps to the species level via PCR of the mitochondrial *cytochrome oxidase subunit I* (*COI*) gene. Annealing temperature (T_A_) and predicted amplicon size in base pairs (bp) following PCR are included.

## 3. Results

### 3.1 Similar diapause phenotypes across wasp species, but different incidence of those phenotypes

Each wasp species exhibited the same three diapause phenotypes (Fig. 1) as their fly host, when non-overwintered individuals collected in 2009 were exposed to constant warm temperature (21°C). Wasps that entered either weak or prolonged diapause showed an initial metabolic rate depression as they entered diapause, and an increase in metabolic rate associated with exit from diapause (Fig. 2). By approximately 20 d post-pupariation, appreciable differences in metabolic rate were evident between non-diapause (high metabolic rate) and weak or prolonged diapause (low metabolic rate) phenotypes (Fig. 2). These non-diapause wasps eclosed as adults in 35 d or less post-pupariation, while weak diapause wasps eclosed between 36 and 100 d post-pupariation, and prolonged diapause wasps did not eclose within 100 d at 21°C (Fig. S1).

**Figure 1.**
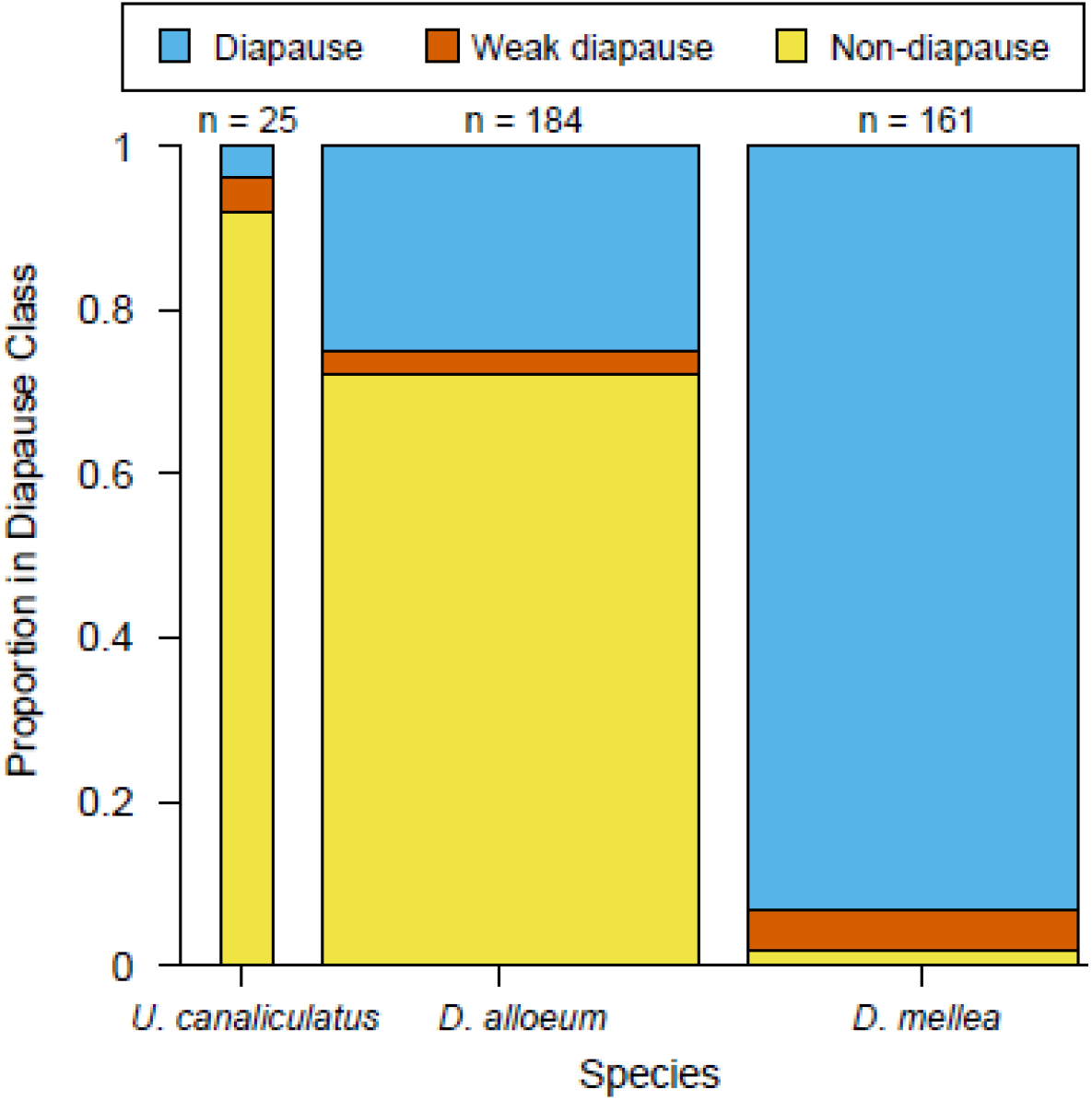
Proportion of non-diapause (yellow), weak diapause (red), and prolonged diapause (blue) phenotypes in *Utetes canaliculatus, Diachasma alloeum*, and *Diachasmimorpha mellea* when exposed to continuous warm conditions (21°C) following fly host pupariation. Bar width is proportional to the number of each wasp species detected from collections made in 2009. The number of individuals of each diapause phenotype are given in Table S1. Data were combined from experiments that used respirometry (Fig. 2) and eclosion time (Fig. S1) to determine diapause phenotype.

**Figure 2.**
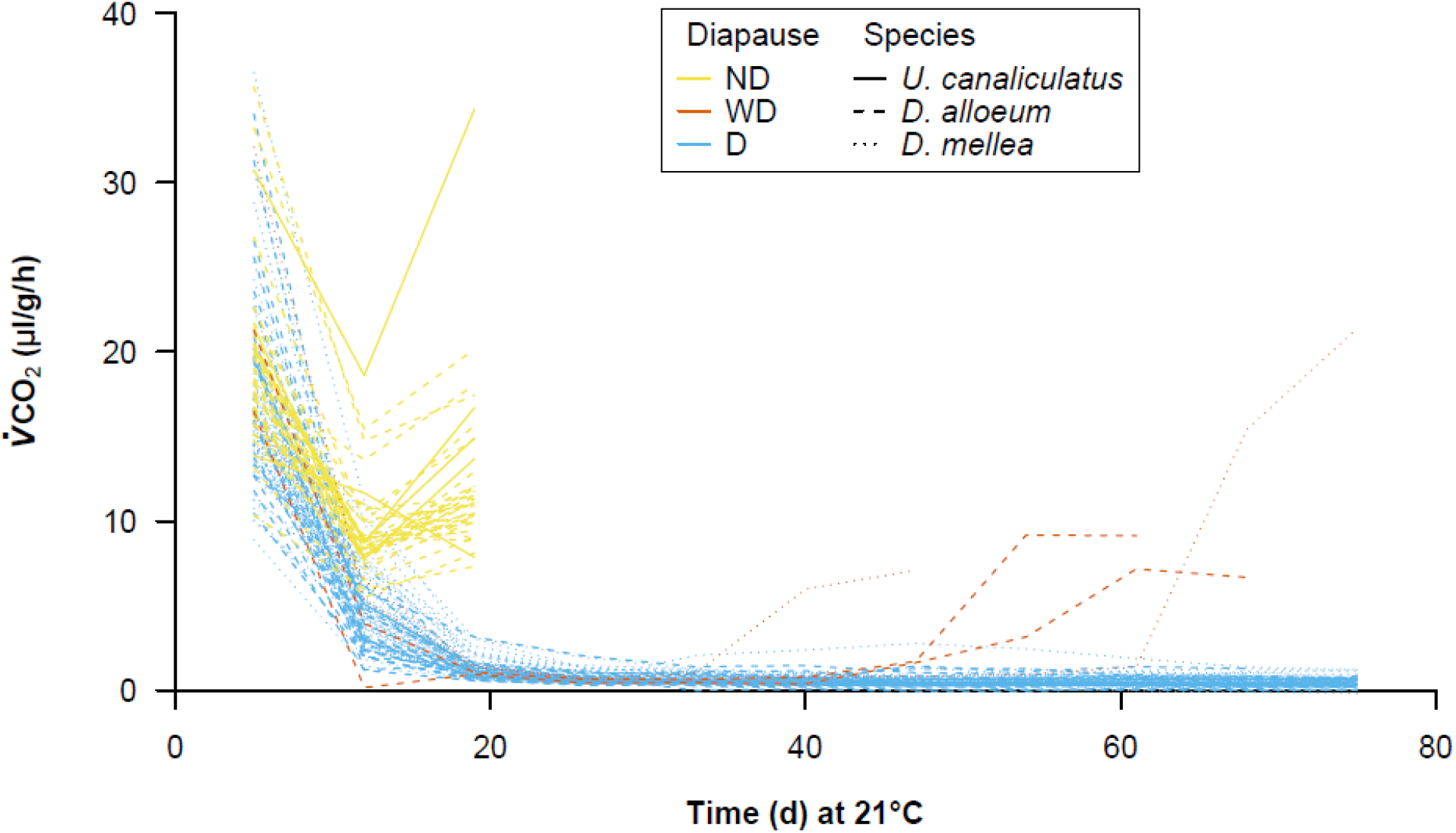
Metabolic rates over time at 21°C of endoparasitoid wasps that never entered diapause (non-diapause; ND = yellow), entered diapause but eclosed within 100 days (weak diapause; WD = red), or entered a prolonged diapause (D = blue). Wasp species can be identified via line type: solid, *Utetes canaliculatus*; dashed, *Diachasma alloeum*; dotted, *Diachasmimorpha mellea*.

Although each wasp species exhibited the same three diapause phenotypes, the proportion of wasps exhibiting each diapause phenotype differed among wasp species (G = 248.8, *P* < 0.001). The weak diapause phenotype occurred in less than 5% of each wasp species (Fig. 1). In contrast, more than 70% of *D. alloeum* and *U. canaliculatus* averted diapause (non-diapause), with less than 25% entering a prolonged diapause (Fig. 1). Conversely, 93% of *D. mellea* entered a prolonged diapause, and only ∼2% displayed the non-diapause phenotype (Fig. 1).

### 3.2 Parasitism rates and relative abundance of parasitoid species

The proportion of fly puparia that contained wasps was similar in our two collections: parasitism rates were 27% and 35% in 2009 and 2019, respectively. However, the relative abundance of each parasitoid wasp species differed between the two collections, which may reflect natural variation between collection years and field sites (Forbes et al., 2010; Hood et al., 2015). In 2009 (Grant, MI), *D. alloeum* was the most abundant species (49%), followed by *D. mellea* (43%) and *U. canaliculatus* (8%). Conversely, in 2019 (East Lansing, MI), the most abundant wasp species was *U. canaliculatus* (67% of the wasps), followed by *D. alloeum* (28%), and *D. mellea* (5%). The egg parasitoid, *U. canaliculatus*, attacks *R. pomonella* earlier in the season than the other two wasp species that attack late instar fly larvae (Forbes et al., 2010; Hood et al., 2015). It is therefore possible that our 2019 field collection occurred prior to much of the natural parasitism of flies by *D. alloeum* and *D. mellea*, resulting in the unusually high relative abundance of *U. canaliculatus*.

### 3.3 Similar acute cold tolerance of D. alloeum, U. canaliculatus, and R. pomonella

Both *U. canaliculatus* and *D. alloeum* had the same cold tolerance strategy as their fly host, *R. pomonella*. After chilling durations of 8 and 16 weeks, these two wasp species and *R. pomonella* survived brief supercooling to low temperatures (c. –20°C) but did not survive internal ice formation (freezing), and were therefore classified as freeze-avoidant (Table 2). Unfortunately, there were not enough *D. mellea* in the 2019 collections (N = 2) to make a conclusion about their cold tolerance strategy. However, *D. mellea* are more likely to be freeze-avoidant than freeze-tolerant as the two individuals that froze did not survive (Table 2). The similarity in cold tolerance strategy between *R. pomonella* and at least two of its parasitoids allows us to compare their tolerance of acute exposure to extreme low temperatures by measuring supercooling point (SCP).

**Table 2.**
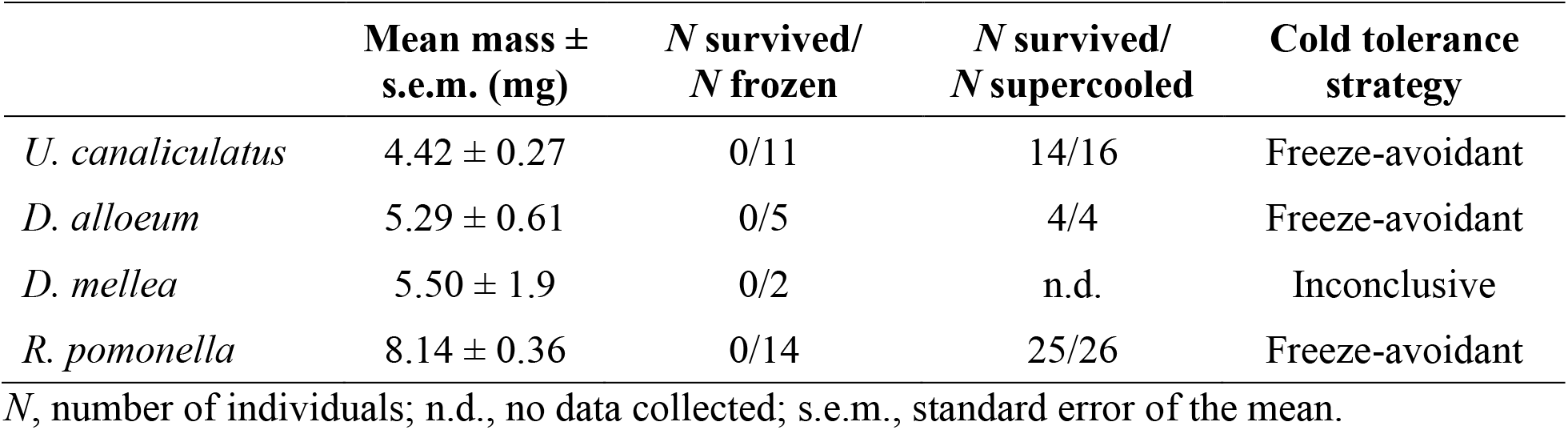
Cold tolerance strategy of endoparasitoid wasps *Utetes canaliculatus, Diachasma alloeum*, and *Diachasmimorpha mellea*, and their fly host *Rhagoletis pomonella*. Data were combined from individuals cooled after 8 and 16 weeks of chilling.

*Utetes canaliculatus, D. alloeum*, and their fly host *R. pomonella* can survive acute exposures to extreme low temperatures similarly well. The temperature at which individuals froze (SCP) was similar among all four species (ANCOVA; Species *F*_1,72_ = 0.315, *P* = 0.815), with most values ranging from –16°C to –22°C (Fig. 3). Therefore, the three freeze-avoidant species (Table 2) can remain unfrozen (supercooled) and alive at similarly low temperatures. In addition, SCP did not vary with insect mass (ANCOVA; Mass *F*_1,68_ = 0.059, *P* = 0.809), or chill duration (ANCOVA; Chill Duration *F*_1,68_ = 0.365, *P* = 0.548). Acute cold tolerance was therefore unaffected by acclimation to prolonged chilling, as SCP did not vary over time.

**Figure 3.**
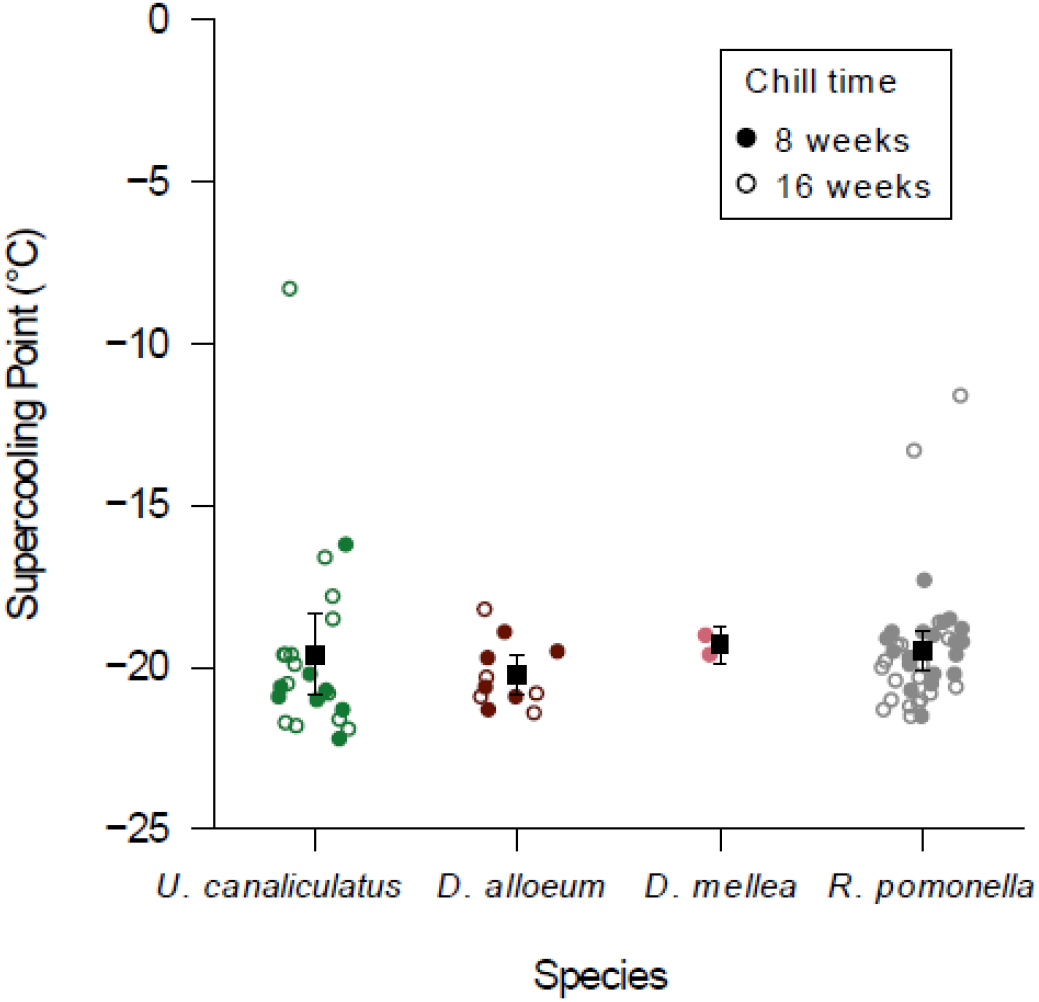
Supercooling point (SCP) temperature of *Utetes canaliculatus* (green), *Diachasma alloeum* (maroon), and *Diachasmimorpha mellea* (pink); and their host *Rhagoletis pomonella* (grey). Square points represent the mean ± 95% confidence interval for each species. Each circular point represents the SCP from one individual measured after 8 weeks (filled) or 16 weeks (open).

*Diachasmiomorpha mellea* (N = 2) froze at similar temperatures to the other species, but further study with more individuals is required to determine their ability to survive these acute low temperatures.

### 3.4 Similar chronic cold tolerance of D. alloeum and U. canaliculatus

While the sample size for *D. mellea* was too low to determine its chronic cold tolerance, we observed similar responses to prolonged chilling in *D. alloeum* and *U. canaliculatus*. The short chilling duration of 8 weeks resulted in wasps eclosing from 35% of fly puparia (114/325; Fig. 4). In contrast, following 16 or 24 weeks of chilling, wasps eclosed from only 8% (27/325) and 1% (3/325) of fly puparia, respectively (Fig. 4), suggesting that prolonged chilling decreased parasitoid survival to adulthood. The proportion of each wasp species that eclosed after chilling did not vary with chill duration (G = 3.322, *P* = 0.506; Fig. 4). Therefore, prolonged chilling caused similar mortality over time in *D. alloeum* and *U. canaliculatus*.

**Figure 4.**
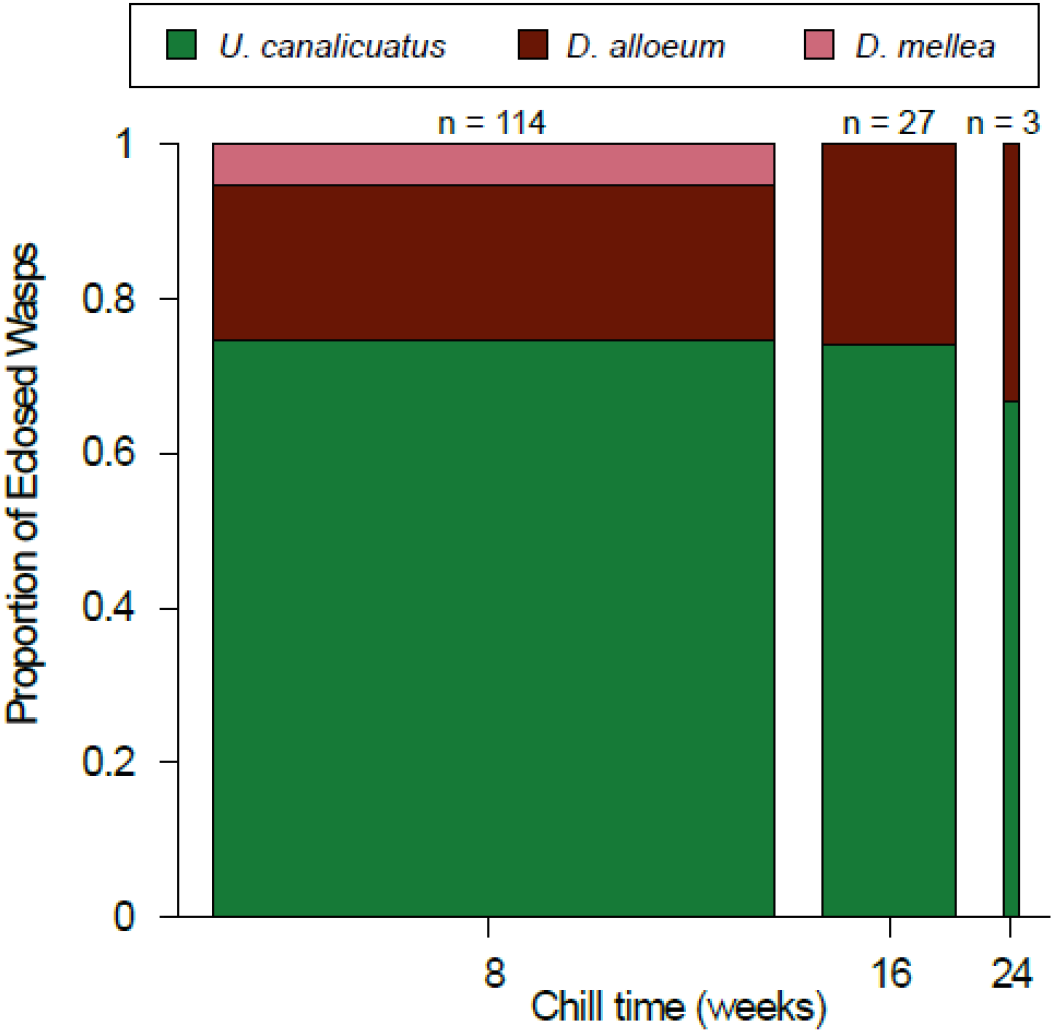
Survival to adulthood of *Utetes canaliculatus* (green), *Diachasma alloeum* (maroon), and *Diachasmimorpha mellea* (pink) after 8, 16, and 24 weeks of chilling at 4°C. Total number of fly pupae, including those parasitized by wasps, exposed to each chill duration was 325. Bar width is proportional to the number of wasps that eclosed; total number eclosed is indicated above each bar. The eclosion time and number of eclosed individuals are available in the Figs. S6 and S7.

## 4. Discussion

### 4.1 Similar diapause phenotypes within and across trophic levels

The three endoparasitoid wasps in our study exhibited the same three diapause phenotypes as their fly host (Calvert et al., 2022; Dambroski and Feder, 2007), suggesting that temperature-sensitive diapause flexibility is conserved within and across trophic levels. In particular, our results suggest that the weak diapause phenotype is more widespread than previously thought (Masaki, 2002; Wilsterman et al., 2021). Although the prolonged laboratory warming treatment under which we observe these phenotypes may not reflect normal natural conditions for these populations, we show that diapause is not obligate for any of the parasitoid wasp species attacking hawthorn-infesting *R. pomonella*, a pattern that is similar for the fly (Calvert et al., 2022; Dambroski and Feder, 2007; Feder et al., 1993; Toxopeus et al., 2021). In *R. pomonella*, there are strong genomic signatures that distinguish weak diapause flies from those that enter prolonged diapause or completely avert diapause (Calvert et al., 2022). In addition, the propensity of *R. pomonella* to avert or prematurely terminate diapause can be influenced by the host-origin of flies (apple or hawthorn), latitude, and the temperatures that they are exposed to during development (Calvert et al., 2022; Dambroski and Feder, 2007). Future work is needed to determine whether similar environmental and genomic factors impact diapause in the guild of parasitoid wasps, which could provide insight the evolution of diapause phenotypes.

In addition to exhibiting the same diapause phenotypes, parasitoids and their fly host had similar metabolic rate trajectories and eclosion timing at warm temperatures. *Rhagoletis pomonella* that enter prolonged or weak diapause suppress their metabolic rate within 10 d post-pupariation, and elevate their metabolic rate at the end of diapause, prior to adult eclosion (Calvert et al., 2022; Powell et al., 2020; Ragland et al., 2009). The weak and prolonged diapause wasps in our study took longer (up to 20 d) to suppress their metabolic rate post-pupariation, but the overall pattern of metabolic rate suppression and elevation at warm temperatures was similar to *R. pomonella*. Eclosion phenology of non-overwintered wasps was also similar to hawthorn-infesting *R. pomonella* from the same population (Calvert et al., 2022). For example, non-diapause wasps eclosed in less than 35 d post-pupariation, later than but overlapping with non-diapause *R. pomonella* that eclose in less than 31 d. In addition, the range of eclosion times for weak diapause *R. pomonella* was similar to that of the wasps, with most eclosing in less than 85 d under prolonged warming. In all four species, individuals that displayed the prolonged diapause phenotype remained uneclosed at warm temperatures for at least 100 d post-pupariation. The impact of diapause phenotype on eclosion phenology is a strong driver of life history timing in *R. pomonella* (Powell et al., 2020) and others species (Hodek and Hodková, 1988; Masaki, 2002; Tauber and Tauber, 1976). If similar mechanisms underlie diapause phenotypes in *R. pomonella* and its parasitoid wasps, this common diapause flexibility may contribute to synchronization of eclosion phenology within and across these trophic levels (Hood et al., 2015).

### 4.2 Different temperature sensitivity of diapause phenotypes within and across trophic levels

Although the three endoparasitoid wasps in our study exhibited the same three diapause phenotypes as their host, the prevalence of each diapause phenotype varied among species. While less than 25% of hawthorn-infesting *R. pomonella* from the same population avert diapause under warm (non-winter) conditions (Calvert et al., 2022), the majority (>70%) of *D. alloeum* and *U. canaliculatus* exhibited the non-diapause phenotype, with few *D. mellea* (< 2%) averting diapause. This high incidence of the non-diapause phenotype in *D. alloeum* and *U. canaliculatus* at 21°C was unexpected, given that diapause can greatly enhance overwintering survival in many insect species (Hahn and Denlinger, 2011; Koštál, 2006; Toxopeus et al., 2021). However, most populations of *U. canaliculatus* and *D. alloeum* attacking hawthorn-infesting flies would rarely experience prolonged pre-winter warming due to the late fruiting phenology of hawthorns (Dambroski and Feder, 2007). Therefore this diapause aversion by *D. alloeum* and *U. canaliculatus* may not often occur in nature.

Although present in all parasitoid species that we studied, weak diapause was rare (< 5%), similar to the low abundance (4%) of this phenotype in their hawthorn-infesting fly host (Calvert et al., 2022). However, weak diapause is not always a rare phenotype; within *R. pomonella*, apple-infesting populations tend to have higher incidence (> 25%) of weak diapause than those that infest hawthorns (Calvert et al. 2022). To determine whether the incidence of weak diapause is generally similar between *R. pomonella* and its parasitoids, future studies should compare incidence of weak diapause in parasitoids from *R. pomonella* populations displaying a range of weak diapause prevalence. In addition, almost all of the 24 species of *Rhagoletis* in North America are attacked by a diverse guild of parasitoids (Forbes et al., 2010; Hood et al., 2015). Members of the same parasitoid guild that attacks *R. pomonella* can also attack the blueberry-infesting *Rhagoletis mendax*, the snowberry-infesting *Rhagoletis zephyria*, the cherry-infesting *Rhagoletis cingulata*, and the undescribed flowering dogwood fly (Forbes et al., 2010; Hood et al., 2015). A comparative approach across these taxa would provide further insight into the factors that influence the prevalence of (weak) diapause in multiple species (Masaki, 2002; Wilsterman et al., 2021).

*Diachasmimorpha mellea* had a slightly higher prevalence (93% vs. 72%) of prolonged diapause compared to their fly host from the same population (Calvert et al., 2022), in contrast to *U. canaliculatus* and *D. alloeum* that tended to avert diapause under prolonged warm conditions. The insensitivity of diapause to warm temperatures in *D. mellea* may be partially explained by differences in geographic distribution among parasitoid species. *Diachasmimorpha mellea* have a relatively southern distribution, and are abundant at these lower latitudes (Forbes et al., 2010; Rull et al., 2009). A higher recalcitrance to temperature-induced aversion of diapause could benefit species that experience a long, warm fall at lower latitudes, ensuring that these individuals remain in diapause until conditions are permissive post-winter (Dambroski and Feder, 2007; Hodek and Hodková, 1988; Wilsterman et al., 2021). If *D. mellea* has invaded northern regions (e.g., our study sites) more recently than the other parasitoid species, most *D. mellea* may still be programmed to enter prolonged diapause under warm conditions.

As fall conditions are predicted to become increasingly warmer with climate change (Le Lann et al., 2021; Marshall et al., 2020), we may observe shifts in the composition of the parasitoid guild that attacks *R. pomonella* due to interspecific differences in temperature-sensitivity of diapause phenotypes. Based on current differences in parasitoid abundance across latitudes, in the future our sites at northern latitudes may resemble southern latitudes under current climate conditions, with a higher abundance of *D. mellea*, and lower abundance of *D. alloeum* and *U. canaliculatus* (Forbes et al. 2010). Understanding the interplay between temperature, flexibility in diapause phenotypes, and life history timing will be critical for predicting these potential shifts in the composition of the parasitoid guild.

### 4.3 Similar acute cold tolerance within and across trophic levels

The similarity in cold tolerance strategy and SCP among *U. canaliculatus, D. alloeum*, and *R. pomonella* suggests limited variation in acute cold tolerance within and across trophic levels in this system. These three species were freeze-avoidant, similar to several other species of parasitoid wasps and their insect hosts that are unable to survive prolonged periods of freezing temperatures (Amiresmaeili et al., 2020; Carrillo et al., 2005; Hanson et al., 2013; Li et al., 2014). This phenotype is perhaps unsurprising, as most insects are either freeze-avoidant or chill-susceptible, and freeze tolerance is a comparatively rare cold tolerance strategy (Overgaard and MacMillan, 2017; Sinclair et al., 2003; Toxopeus and Sinclair, 2018). In addition, the cold tolerance we observed may not be strongly impacted by the host fruit environment of *R. pomonella*. The hawthorn-infesting flies in this study had the same cold tolerance strategy (freeze avoidance) and froze at similar temperatures (c. -20°C) to *R. pomonella* that infest apples (Toxopeus et al., 2021). Future work is needed to determine whether similar physiological mechanisms underlie the similarity in acute cold tolerance among these closely-interacting species.

The low interspecific variation in acute cold tolerance among *R. pomonella* and its parasitoids may be due to the rarity with which these insects experience acute exposures to extreme low temperatures. The low temperatures we used in the acute cold tolerance experiments (c. -20°C) rarely occur in the overwintering habitat of *R. pomonella*: the soil in which fly pupae overwinter is unlikely to decrease below -5°C, especially if snow cover is present (Filchak et al., 2000). All three endoparasitoid wasps should experience a similar winter microclimate, including any protection conferred by the fly puparium that may, for example, minimize contact with ice in the soil that could initiate freezing. Tolerance of freezing or acute exposure to extreme low temperatures therefore likely has a limited impact on the overwintering ability of the species in this study.

### 4.4 Similar chronic cold tolerance within a trophic level

The two most abundant wasps in our study – *U. canaliculatus* and *D. alloeum –* exhibited similar survival following a simulated winter of chilling at 4°C, further supporting a lack of interspecific variation in cold tolerance in this system. The similarity in chronic cold tolerance of *D. alloeum* and *U. canaliculatus* could be influenced by similarities in wasp diapause status or by host physiology. For example, the greater tendency of *D. alloeum* and *U. canaliculatus* to avert diapause may result in poor energy reserve accumulation (Hahn and Denlinger, 2011) and higher mortality due to energy drain at mild winter temperatures. Conversely, characteristics of host physiology, such as diet or whether the host is in diapause, can have strong effects on parasitoid cold tolerance physiology (Li et al., 2015; Tougeron et al., 2019; but see Alford et al., 2017). Given that all the wasps in our cold tolerance experiments were collected from the same population of hawthorn-infesting *R. pomonella*, many of the wasps likely had access to similar host-related resources, resulting in similar overwintering success (Hahn and Denlinger, 2011; Toxopeus and Sinclair, 2018). However, even within a single population of flies, each parasitoid species may have different access to energy reserves due to multiple wasps parasitizing a single host (Hood, 2016; Hood et al., 2021, 2012), variation in energy reserve accumulation among hosts prior to pupariation (Li et al., 2014), or differential use of host resources by the parasitoids (Le Lann et al., 2012). Additional work is needed to determine whether diapause phenotypes interact with chronic cold tolerance and, for example, if species with higher incidence of prolonged diapause such as *D. mellea* are more likely to increase in abundance with changing climates.

Overall, chronic cold tolerance of the endoparasitoid wasps in our study was similar to or lower than previous work on these species. Across a 10-year period, parasitoid wasps eclosed post-chill from 20-28% of *R. pomonella* puparia collected from five sites in the midwestern USA, including sites used in this study (Hood, 2016; Hood et al., 2015). The chilling periods in these past studies ranged from 16 – 32 weeks (Hood, 2016; Hood et al., 2015). We saw higher prevalence of eclosed wasps (35%) in our study following 8 weeks of chilling, but much lower prevalence of eclosed wasps (8% and 1%) when our chilling durations were comparable to these previous studies (16 and 24 weeks, respectively). We expected the wasps to be well-adapted to 16 weeks of chilling at 4°C, because many overwintering sites are typically at or below this temperature for 4 months (National Climatic Data Center; NCDC). The high mortality we observed may be due to variation in parasitoid survival across collection sites and years (Hood et al., 2015) or differences in our 2019 laboratory set-up compared to other studies. For example, the 10 d pre-winter period prior to chilling was at a higher temperature (25°C) than previous studies (summarized in Hood et al., 2015), which may contribute to the draining of energy reserves prior to chilling (Hahn and Denlinger, 2011). In addition, most other *Rhagoletis*-parasitoid studies have overwintered individuals on moist vermiculite rather than empty Petri plates (Dambroski and Feder, 2007; Hood et al., 2015; Rull et al., 2009); therefore the parasitoids in our study may have been exposed to different levels of desiccation stress during chilling. Future work is required to examine whether multiple stressors, humidity and pre-chill temperature exposure, affect the susceptibility of parasitoid wasps to chilling-induced mortality from chronic cold injury (Overgaard and MacMillan, 2017) or energy drain (Hahn and Denlinger, 2011) during chilling.

### 4.5 Conclusions

In this study, we observed several similarities between the endoparasitoid wasps of *R. pomonella* and their host, suggesting conservation of diapause and cold tolerance strategies within the parasitoid guild and across trophic levels. The same three diapause phenotypes (non-diapause, weak diapause, and prolonged diapause) exhibited by *R. pomonella* were present in the three wasp species we studied. Although much work has been done on diapause in *R. pomonella* (Calvert et al., 2022; Doellman et al., 2019; Dowle et al., 2020; Egan et al., 2015; Hood et al., 2020; Powell et al., 2020; Ragland et al., 2017), the diapause physiology of the guild of endoparasitoids that attack the fly is comparatively understudied (Hood et al., 2015). Future work should focus on whether flexibility in diapause phenotypes of the parasitoids facilitates life history synchronization of these closely-interacting wasp and fly species. The two most abundant wasp species, *D. alloeum* and *U. canaliculatus*, had remarkably similar acute and chronic cold tolerance. More work needs to be done to determine whether the rarer parasitoid, *D. mellea*, can survive prolonged chilling due to its very high incidence of prolonged diapause. The interaction between warmer and more prolonged pre-winter conditions due to climate change (Le Lann et al., 2021; Marshall et al., 2020), incidence of different diapause phenotypes, and overwintering success will be important for understanding this community of insects and the plants they infest.

## Acknowledgements

We thank James Smith and Thomas Powell for helping collect infested fruit and Lahari Gadhey, Isaiah Sower, Matin Sanaei, and Joseph Tucker for collecting pupae and helping track post-chill eclosion.

## Declarations of interest

None. The authors declare that they have no competing interests.

## Funding

This work was funded by National Science Foundation IOS 1700773 and DEB 1638951 grants to GJR, a Natural Sciences and Engineering Research Council of Canada (NSERC) Discovery Grant to JT, a BioTalent Canada and St. Francis Xavier University Biology Chiasson Award to TM, a University of Colorado, Denver Undergraduate Research Opportunity Program grant to LA, an Indiana Academy of Science, the Entomological Society of America, and Sigma Xi awards to GRH, National Science Foundation grants IOS 1257298 and DEB 1639005 as well as Florida Agricultural Experiment Station Hatch Project FLA-ENY-005943 to DAH.

## Data availability

All data and R code for statistical analysis are available at https://github.com/jtoxopeus/parasitoid-coldtolerance.

## Notes

https://github.com/jtoxopeus/parasitoid-coldtolerance

### Competing Interest Statement

The authors have declared no competing interest.

