## Supplementary Figures and Tables for "Cold tolerance and diapause within and across trophic levels: endoparasitic wasps and their fly host have similar phenotypes"

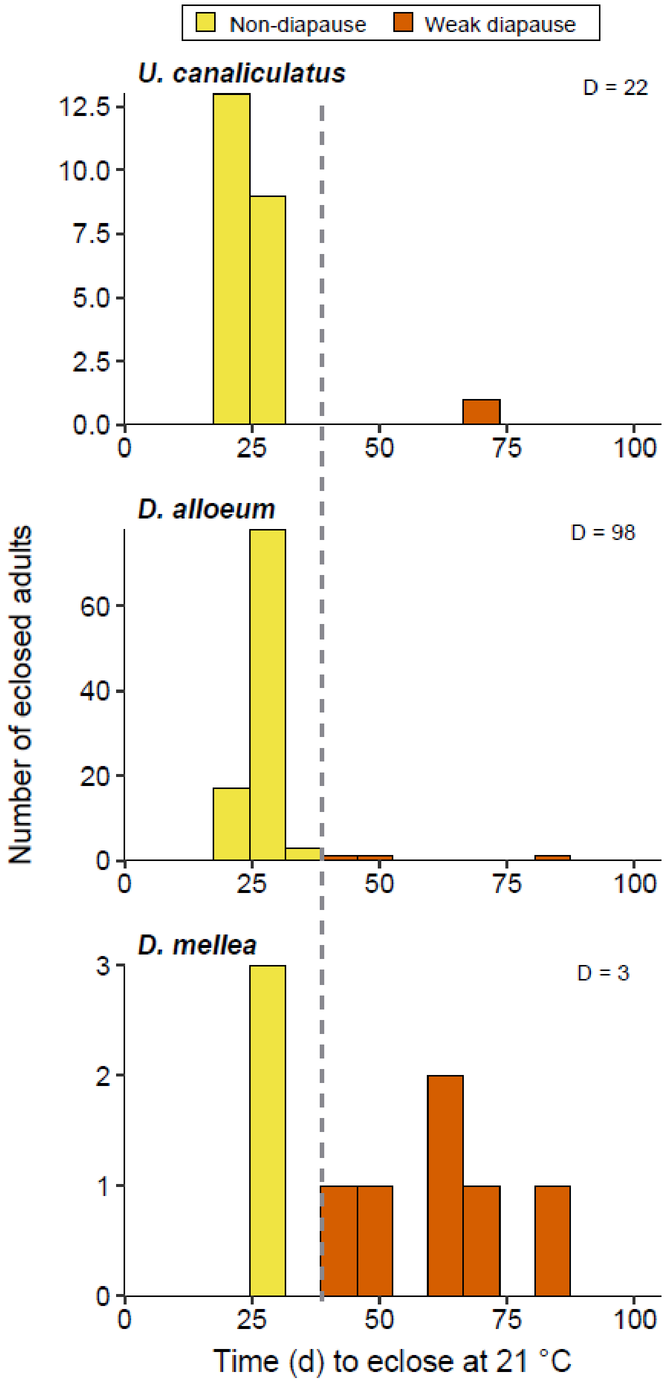


**Figure S1.** Distribution of time to eclosion at 21°C for **(A)** *Utetes canaliculatus*, **(B)** *Diachasma alloeum*, and **(C)** *Diachasmimorpha mellea* in the absence of chilling. Each bar represents the total number of individuals for a particular species that eclosed in a 7-day period. Individuals to the left of the dashed line (early eclosing phenotype) were classified as non-diapause (ND); individuals to the right of the dashed line (late eclosing phenotype) were classified as weak diapause (WD). The number of wasps that remained uneclosed after 100 days (prolonged diapause, D) is indicated in the top right of each panel.

**Table S1**. Comparison of the proportion ± 95% confidence interval and number (N) of pupae in each diapause class when diapause phenotype is determined via respirometry or eclosion time at 21°C. The 95% confidence intervals overlap for proportions calculated from either experiment (respirometry or eclosion time), therefore we pooled the data shown here for display in Fig. 1.

| **Species** | **Phenotyping experiment** | **Proportion in each diapause class (N)** | | |
|---|---|---|---|---|
| | | Non-diapause | Weak diapause | Diapause |
| *Diachasma alloeum* | Respirometry | 0.58±0.16 (35) | 0.03±0.25 (2) | 0.38±0.20 (23) |
| | Eclosion time | 0.79±0.08 (98) | 0.02±0.17 (3) | 0.19± 0.16 (23) |
| *Diachasmimorpha mellea* | Respirometry | 0 (0) | 0.03±0.25 (2) | 0.97±0.05 (59) |
| | Eclosion time | 0.03±0.19 (3) | 0.06±0.19 (6) | 0.91±0.06 (91) |
| *Utetes canaliculatus* | Respirometry | 0.86±0.28 (6) | 0 (0) | 0.14±0.69 (1) |
| | Eclosion time | 0.96±0.09 (22) | 0.04±0.4 (1) | 0 (0) |

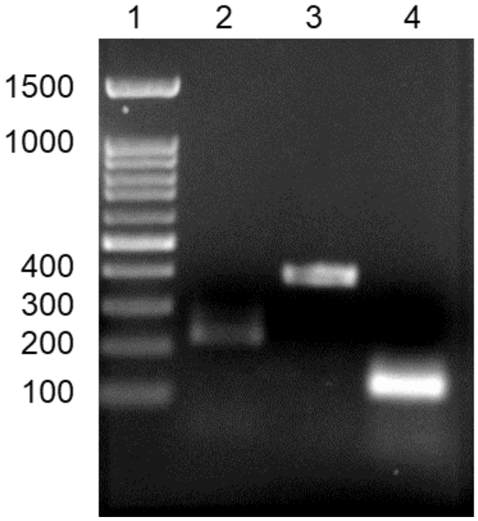


**Figure S2.** Representative agarose gel of PCR products of a portion of the *cytochrome oxidase I* (*COI*) mitochondrial gene with the species-specific primers in Table 1. Each primer pair was used to amplify a product from DNA of one adult of the appropriate species (identified by morphology) using the PCR conditions described in the main text. Lanes: 1, ladder, with relevant sizes indicated to the left in bp (base pairs); 2, *Utetes canaliculatus*; 3, *Diachasma alloeum*; 4, *Diachasmimorpha mellea*.

```
EU881544.1      ------------------------------------------------TGAGCTGGGGTG    12
D.alloeum       ------------------------------------------------------------    0
KR890748.1      ------------------------GTTTTGTATTTTTTGTATGGGATTTGAGCTGGGGTG    36
KU511212.1      TCAACAAATCATAAAGATATTGGTGTTTTGTATTTTTTGTATGGGATTTGAGCTGGGGTG    60


EU881544.1      TTAGGTTTATCTATAAGAATAATTATTCGATTAGAATTAGGGATACCTGGGAGATTGTTA    72
D.alloeum       ------------------------------------------------------------    0
KR890748.1      TTAGGTTTATCTATAAGAATAATTATTCGATTAGAATTAGGGATACCTGGGAGATTGTTA    96
KU511212.1      TTAGGTTTATCTATAAGAATAATTATTCGATTAGAATTAGGGATACCTGGGAGATTGTTA    120


EU881544.1      AATGATCAGATTTATAATAGAATAGTAACTGCTCATGCTTTTGTTATAATTTTTTTTACA    132
D.alloeum       ---------------------------------------TTTGTTATAATTTTTTTTACA    21
KR890748.1      AATGATCAGATTTATAATAGAATAGTAACTGCTCATGCTTTTGTTATAATTTTTTTTACA    156
KU511212.1      AATGATCAGATTTATAATAGAATAGTAACTGCTCATGCTTTTGTTATAATTTTTTTTACA    180
                                                       *********************

EU881544.1      GTTATACCTATTATAATTGGGGGGTTTGGGAATTGATTAATTCCTTTAATATTAGGGGTT    192
D.alloeum       GTTATACCTATTATAATTGGAGGGTTTGGGAATTGATTAATTCCTTTAATATTAGGGGTT    81
KR890748.1      GTTATACCTATTATAATTGGAGGGTTTGGGAATTGATTAATTCCTTTAATATTAGGGGTT    216
KU511212.1      GTTATACCTATTATAATTGGAGGGTTTGGGAATTGATTAATTCCTTTAATATTAGGGGTT    240
                ******************** ***************************************

EU881544.1      CCTGATATAGCCTTTCCTCGAATAAATAATATAAGATTTTGATTATTGAGACCTTCTATA    252
D.alloeum       CCTGATATAGCCTTTCCTCGAATAAATAATATAAGATTTTGATTATTGAGACCTTCTATA    141
KR890748.1      CCTGATATAGCCTTTCCTCGAATAAATAATATAAGATTTTGATTATTGAGACCTTCTATA    276
KU511212.1      CCTGATATAGCCTTTCCTCGAATAAATAATATAAGATTTTGATTATTGAGACCTTCTATA    300
                ************************************************************

EU881544.1      ATTTTGTTAATATTGAGAATATTATTAAATTTAGGTGCTGGAACTGGTTGGACAATTTAT    312
D.alloeum       ATTTTGTTAATATTGAGAATATTATTAAATTTAGGTGCTGGAACTGGTTGGACAATTTAT    201
KR890748.1      ATTTTGTTAATATTGAGAATATTATTAAATTTAGGTGCTGGAACTGGTTGGACAATTTAT    336
KU511212.1      ATTTTGTTAATATTGAGAATATTATTAAATTTAGGTGCTGGAACTGGTTGGACAATTTAT    360
                ************************************************************

EU881544.1      CCTCCTTTATCATCTAGATTAGGTCATAGGGGGTTAGCAGTAGATTTATTAATTTTTAGT    372
D.alloeum       CCTCCTTTATCATCTAGATTAGGTCATAGGGGGTTAGCAGTAGATTTATTAATTTTTAGT    261
KR890748.1      CCTCCTTTATCATCTAGATTAGGTCATAGGGGGTTAGCAGTAGATTTATTAATTTTTAGT    396
KU511212.1      CCTCCTTTATCATCTAGATTAGGTCATAGGGGGTTAGCAGTAGATTTATTAATTTTTAGT    420
                ************************************************************

EU881544.1      TTACATTTAGCTGGGGTATCATCAATTATGGGGGCAATTAATTTTATTTGTACAATTTTA    432
D.alloeum       TTACATTTAGCTGGGGTATCATCAATTATGGGGGCAATTAATTTTATTTGTACAATTTTA    321
KR890748.1      TTACATTTAGCTGGGGTATCATCAATTATGGGGGCAATTAATTTTATTTGTACAATTTTA    456
KU511212.1      TTACATTTAGCTGGGGTATCATCAATTATGGGGGCAATTAATTTTATTTGTACAATTTTA    480
                ************************************************************

EU881544.1      AATATAAAGCTTTTCATAAAGTTTGAGCAATTAAGTTTATTTATTTGGTCAATTTTAATT    492
D.alloeum       AATATAAAGCTTTTCATAAAGTTTGAGCAATTAA--------------------------    355
KR890748.1      AATATAAAGCTTTTCATAAAGTTTGAGCAATTAAGTTTATTTATTTGGTCAATTTTAATT    516
KU511212.1      AATATAAAGCTTTTCATAAAGTTTGAGCAATTAAGTTTATTTATTTGGTCAATTTTAATT    540
                **********************************

EU881544.1      ACAGCTATTTTATTATTATTATCTTTACCGGTTTTAGCTGGAGCTATTACAATATTATTA    552
D.alloeum       ------------------------------------------------------------    355
KR890748.1      ACAGCTATTTTATTATTATTATCTTTACCGGTTTTAGCTGGAGCTATTACAATA------    570
KU511212.1      ACAGCTATTTTATTATTATTATCTTTACCGGTTTTAGCTGGAGCTATTACAATATTATTA    600
```

**Figure S3.** Alignment of the positive control *Diachasma alloeum* PCR amplicon to *cytochrome oxidase I* (*COI*) gene sequences from *D. alloeum* vouchers in NCBI. Default parameters were used in Clustal Omega (https://www.ebi.ac.uk/Tools/msa/clustalo/) for alignment Asterisks (*) indicate exact matches between our sequence (*D. alloeum*) and the three NCBI sequences (identified by NCBI accession numbers).

```
KT761492.1      TCAATAAGAATAATTATTCGGATAGAATTAGGGGTCCCTGGTAGGATATTGATAAGAGAT 60
KT761495.1      TCAATAAGAATAATTATTCGGATAGAATTAGGGGTTCCTGGTAGGATATTGATAAGAGAT 60
D.mellea        ------------------------------------------------------------ 0
KT761488.1      TCAATAAGAATAATTATTCGGATAGAATTAGGGGTTCCTGGTAGGATATTGATAAGAGAT 60


KT761492.1      CAACTTTATAATAGTATAGTTACTTCTCATGCTTTTGTTATAATTTTTTTTATAGTTATA 120
KT761495.1      CAACTTTATAATAGTATAGTTACTTCTCATGCTTTTGTTATAATTTTTTTTATAGTTATA 120
D.mellea        -------------------TTACTTCTCATGCTTTTGTTATAATTTTTTTTATAGTTATA 41
KT761488.1      CAACTTTATAATAGTATAGTTACTTCTCATGCTTTTGTTATAATTTTTTTTATAGTTATA 120
                                   *****************************************

KT761492.1      CCTATTATAATTGGTGGGTTTGGTAATTGGTTGGTACCTTTAATATTAGGGGCTCCTGAT 180
KT761495.1      CCTATTATAATTGGTGGGTTTGGTAATTGGTTGGTACCTTTAATATTAGGGGCTCCCGAT 180
D.mellea        CCTATTATAATTGGTGGGTTTGGTAATTGGTTGGTACCTTTAATATTA------------ 89
KT761488.1      CCTATTATAATTGGTGGGTTTGGTAATTGGTTGGTACCTTTAATATTAGGGGCTCCTGAT 180
                ************************************************

KT761492.1      ATAGCTTTCCCTCGAATAAATAATATAAGGTTTTGATTACTTGTTCCTTCTTTATTTTTA 240
KT761495.1      ATAGCTTTCCCTCGAATAAATAATATAAGGTTTTGATTACTTGTTCCTTCTTTATTTTTA 240
D.mellea        ------------------------------------------------------------ 89
KT761488.1      ATAGCTTTCCCTCGAATAAATAATATAAGGTTTTGATTACTTGTTCCTTCTTTATTTTTA 240


KT761492.1      TTAATGTTGAGAAGATTATTAAATTTGGGGGTTGGAACTGGTTGAACAGTTTATCCTCCA 300
KT761495.1      TTAATGTTGAGAAGATTATTAAATTTGGGGGTTGGAACTGGTTGAACAGTTTATCCTCCA 300
D.mellea        ------------------------------------------------------------ 89
KT761488.1      TTAATGTTGAGAAGATTATTAAATTTGGGGGTTGGAACTGGTTGAACAGTTTATCCTCCA 300


KT761492.1      TTGTCATCAAATTTGGGGCATGTAGGTTCATCCGTTGATTTAGCTATTTTTTCTTTACAT 360
KT761495.1      TTGTCATCAAATTTGGGGCATGTAGGTTCATCCGTTGATTTAGCTATTTTTTCTTTACAT 360
D.mellea        ------------------------------------------------------------ 89
KT761488.1      TTGTCATCAAATTTGGGGCATGTAGGTTCATCCGTTGATTTAGCTATTTTTTCTTTACAT 360


KT761492.1      TTGGCGGGAGTTTCTTCAATTATAGGGGCTATTAATTTTATTAGAACAATTTTGAATATA 420
KT761495.1      TTGGCGGGAGTTTCTTCAATTATAGGGGCTATTAATTTTATTAGAACAATTTTGAATATA 420
D.mellea        ------------------------------------------------------------ 89
KT761488.1      TTGGCGGGAGTTTCCTCAATTATAGGGGCTATTAATTTTATTAGAACAATTTTGAATATA 420


KT761492.1      AATTTTTATATAATTAAATTAGATCAGTTAAGTTTATTAATTTGGTCAATTTTAATTACG 480
KT761495.1      AATTTTTATATAATTAAATTAGATCAGTTAAGTTTATTAATTTGGTCAATTTTAATTACG 480
D.mellea        ------------------------------------------------------------ 89
KT761488.1      AATTTTTATATAATTAAATTAGATCAGTTAAGTTTATTAATTTGGTCAATTTTAATTACG 480


KT761492.1      GCTATTTTATTGTTGTTATCTTTGCCTGTTTTAGCTGGTGCTATTACTATGTTGTTAACT 540
KT761495.1      GCTATTTTATTGTTGTTATCTTTGCCTGTTTTAGCTGGTGCTATTACTATGTTGTTAACT 540
D.mellea        ------------------------------------------------------------ 89
KT761488.1      GCTATTTTATTGTTGTTATCTTTGCCTGTTTTAGCTGGTGCTATTACTATGTTGTTAACT 540


KT761492.1      GATCGAAATTTAAATACG    558
KT761495.1      GATCGAAATTTAAATACG    558
D.mellea        ------------------    89
KT761488.1      GATCGAAATTTAAATACG    558
```

**Figure S4.** Alignment of the positive control *Diachasmimorpha mellea* PCR amplicon to *cytochrome oxidase I* (*COI*) gene sequences from *D. mellea* vouchers in NCBI. Default parameters were used in Clustal Omega (https://www.ebi.ac.uk/Tools/msa/clustalo/) for alignment Asterisks (*) indicate exact matches between our sequence (*D. mellea*) and the three NCBI sequences (identified by NCBI accession numbers).

```
KU511220.1          TCTGGTATAGTAGGTTTATCAATAAGATTAATTATTCGTATAGAATTAGGTGTTCCTGGA     60
KT761328.1          ------------------------------------------------------CCTGGA     6
U.canaliculatus     ------------------------------------------------------------     0
KT761344.1          ------------------------------------------------------CCTGGA     6


KU511220.1          AGATTATTAATAAATGATCAAATTTATAATAGAATAGTTACGGCACATGCTTTTGTAATA     120
KT761328.1          AGATTATTAATAAATGATCAAATTTATAATAGAATAGTTACGGCACATGCTTTTGTAATA     66
U.canaliculatus     ------------------------------------------------------------     0
KT761344.1          AGATTATTAATAAATGATCAAATTTATAATAGAATAGTTACGGCACATGCTTTTGTAATA     66


KU511220.1          ATTTTTTTTATAGTTATGCCAATTATAATTGGTGGTTTTGGTAATTGATTAATTCCTTTA     180
KT761328.1          ATTTTTTTTATAGTTATGCCAATTATAATTGGTGGTTTTGGTAATTGATTAATTCCTTTA     126
U.canaliculatus     ------------------------------------------------------------     0
KT761344.1          ATTTTTTTTATAGTTATGCCAATTATAATTGGTGGTTTTGGTAATTGATTAATTCCTTTA     126


KU511220.1          ATATTAGGAGCTCCTGATATAGCTTTCCCTCGTATAAATAATATAAGTTTTTGATTATTA     240
KT761328.1          ATATTAGGAGCTCCTGATATAGCTTTCCCTCGTATAAATAATATAAGTTTTTGATTATTA     186
U.canaliculatus     ------------------------------------------------------------     0
KT761344.1          ATATTAGGAGCTCCTGATATAGCTTTCCCTCGTATAAATAATATAAGTTTTTGATTATTA     186


KU511220.1          ATTCCTTCATTAATATTATTAATTTTGAGAAGATTATTAAATGTTGGTGTTGGTACTGGT     300
KT761328.1          ATTCCTTCATTAATATTATTAATTTTGAGAAGATTATTAAATGTTGGTGTTGGTACTGGT     246
U.canaliculatus     -----------------------------------------Tgttggtgttggtactggt     19
KT761344.1          ATTCCTTCATTAATATTATTAATTTTGAGAAGATTATTAAATGTTGGTGTTGGTACTGGT     246
                                                             *******************

KU511220.1          TGAACAGTTTATCCACCTTTATCTTCAACATTAGGTCATGGTGGATTATCTGTTGATTTA     360
KT761328.1          TGAACAGTTTATCCACCTTTATCTTCAACATTAGGTCATGGTGGGTTATCTGTTGATTTA     306
U.canaliculatus     tgaacagtttatccacctttatcttcaacattaggtcatggtgggttatctgttgattta     79
KT761344.1          TGAACAGTTTATCCACCTTTATCTTCAACATTAGGTCATGGTGGGTTATCTGTTGATTTA     306
                    ******************************************** ***************

KU511220.1          GCTATTTTTTCTTTACATTTAGCAGGTGTTTCTTCAATTATAGGGGCTATTAATTTTATT     420
KT761328.1          GCTATTTTTTCTTTACATTTAGCAGGTGTTTCTTCAATTATAGGGGCTATTAATTTTATT     366
U.canaliculatus     gctattttttctttacatttagcaggtgtttcttcaattataggggctattaattttatt     139
KT761344.1          GCTATTTTTTCTTTACATTTAGCAGGTGTTTCTTCAATTATAGGGGCTATTAATTTTATT     366
                    ************************************************************

KU511220.1          ACTACTATTTTTAATATAAATTTTTTTATAATTAAGTTAGATCAATTAAGATTATTAATT     480
KT761328.1          ACTACTATTTTTAATATAAATTTTTTTATAATTAAGTTAGATCAATTAAGATTATTAATT     426
U.canaliculatus     actactatttttaatataaatttttttataattaagttagatcaatt-------------     186
KT761344.1          ACTACTATTTTTAATATAAATTTTTTTATAATTAAGTTAGATCAATTAAGATTATTAATT     426
                    ***********************************************

KU511220.1          TGATCAATTTTAATTACAGCTATTTTATTATTATTATCTTTACCAGTATTAGCTGGAGCT     540
KT761328.1          TGATCAATTTTAATTACAGCTATTTTATTATTATTATCTTTACCAGTATTAGCTGGAGCT     486
U.canaliculatus     ------------------------------------------------------------     186
KT761344.1          TGATCAATTTTAATTACAGCTATTTTATTATTATTATCTTTACCAGTATTAGCTGGAGCT     486


KU511220.1          ATTACGATATTATTAACTGATCGAAATTTAAATACATCATTTTTTGATTTTTCTGGTGGG     600
KT761328.1          ATTACGATATTATTAACTGATCGAAATTTAAATACATCATTTTTTGATTTTTCTGGT---     543
U.canaliculatus     ------------------------------------------------------------     186
KT761344.1          ATTACTATATTATTAACTGATCGAAATTTAAATACATCATTTTTTGATTTTTCTGGT---     543
```

**Figure S5.** Alignment of the positive control *Utetes canaliculatus* PCR amplicon to *cytochrome oxidase I* (*COI*) gene sequences from *U. canaliculatus* vouchers in NCBI. Default parameters were used in Clustal Omega (https://www.ebi.ac.uk/Tools/msa/clustalo/) for alignment Asterisks (*) indicate exact matches between our sequence (*U. canaliculatus*) and the three NCBI sequences (identified by NCBI accession numbers).

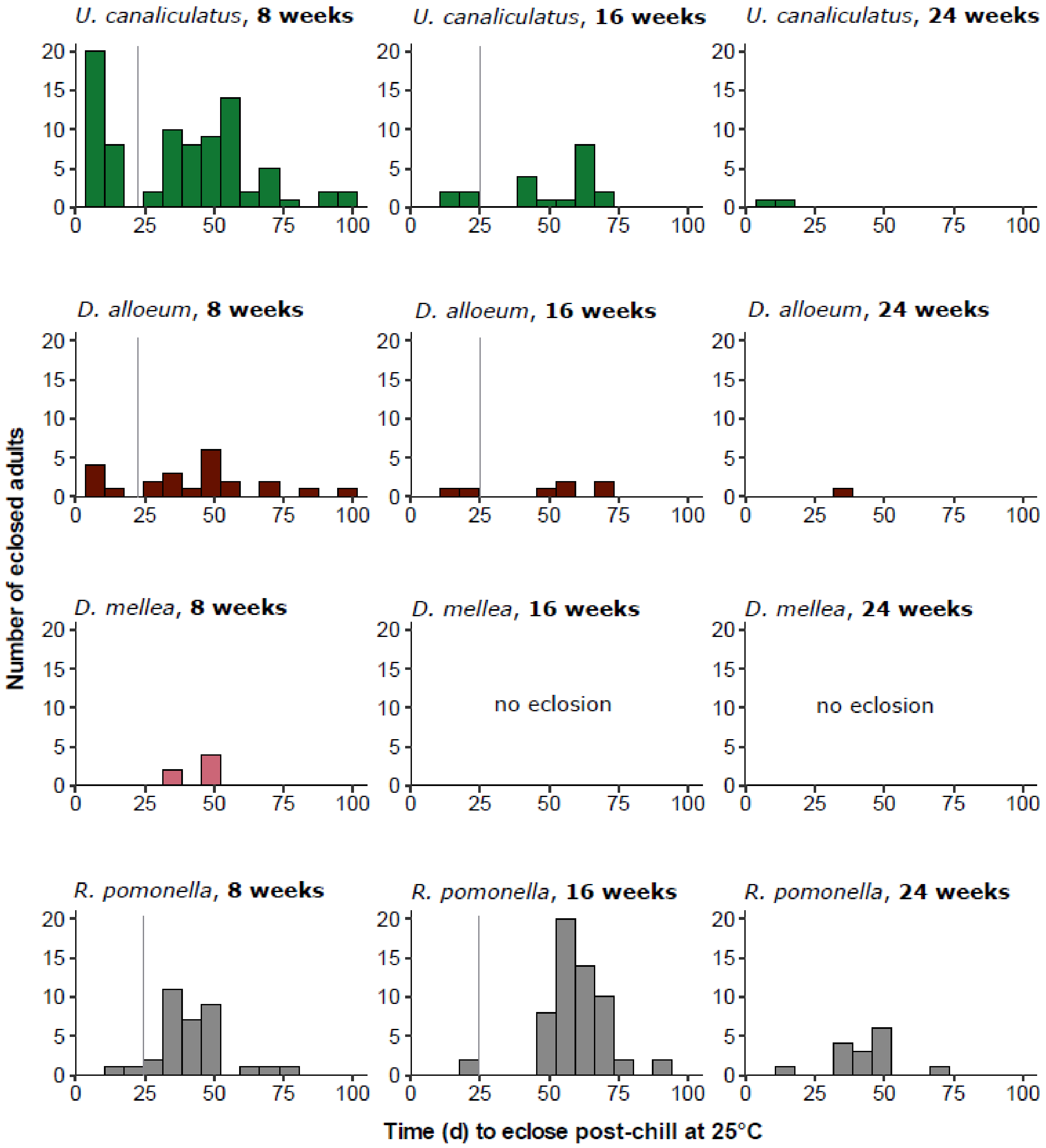


**Figure S6.** Distribution of eclosion phenology for *Rhagoletis pomonella* and the parasitoids that attack the fly (*Diachasma alloeum, Diachasmimorpha mellea, Utetes canaliculatus)* at 25°C after 8, 16, or 24 weeks of chilling at 4°C. For each chill treatment (column), 325 pupae (parasitized and unparasitized) were chilled. Each bar represents the total number of individuals for a particular species that eclosed in a 7-day period (1 week). Individuals with eclosion times less than (to the left of) the vertical grey line likely never entered diapause (non-diapause).

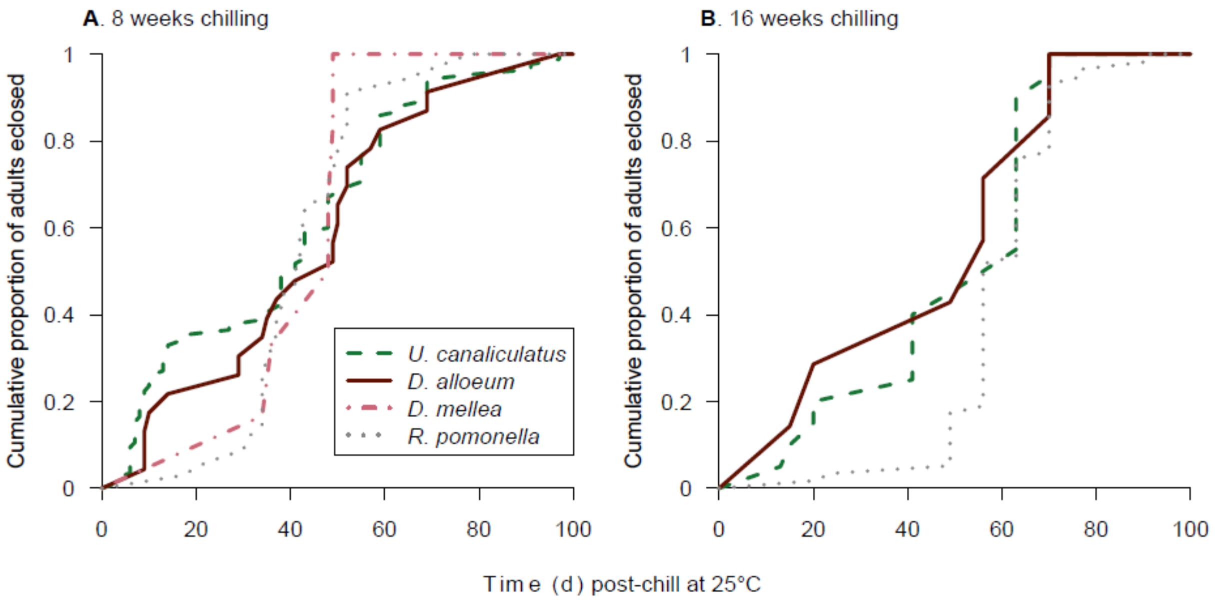


**Figure S7**. Cumulative number of adult flies and parasitoid species that eclosed within 100 days at 25°C after chilling at 4°C for **(A)** 8 weeks or **(B)** 16 weeks. These data are the same as in Fig. S6, but expressed as proportions rather than total numbers of eclosed individuals. Pairwise differences in eclosion times were assessed via Kolmogorov-Smirnov (K-S) tests. The K-S tests revealed that *U. canaliculatus* and *R. pomonella* had different eclosion distributions from each other after 8 weeks ($D$ = 0.329, $P$ = 0.011) and 16 weeks chilling ($D$ = 0.366, $P$ = 0.038). No other pairwise comparisons were significant ($P$ > 0.05). Insufficient wasps eclosed after 24 weeks chilling (< 2 per wasp species) to plot proportion eclosion. However, all data including the 24 week treatment are displayed in Fig. S6.
